# Enhanced top-down sensorimotor processing in somatic anxiety

**DOI:** 10.1101/2021.12.02.471029

**Authors:** Ismail Bouziane, Moumita Das, Karl J Friston, Cesar Caballero-Gaudes, Dipanjan Ray

## Abstract

Functional neuroimaging research on anxiety has traditionally focused on brain networks associated with the psychological aspects of anxiety. Here, instead, we target the somatic aspects of anxiety. Motivated by the growing appreciation that top-down cortical processing plays a crucial role in perception and action, we used resting-state functional MRI data from the Human Connectome Project and Dynamic Causal Modeling (DCM) to characterize effective connectivity among hierarchically organized regions in the exteroceptive, interoceptive, and motor cortices. In people with high (fear-related) somatic arousal, top-down effective connectivity was enhanced in all three networks: an observation that corroborates well with the phenomenology of anxiety. The anxiety-associated changes in connectivity were sufficiently reliable to predict whether a new participant has mild or severe somatic anxiety. Interestingly, the increase in top-down connections to sensorimotor cortex were not associated with fear affect scores, thus establishing the (relative) dissociation between somatic and cognitive dimensions of anxiety. Overall, enhanced top-down effective connectivity in sensorimotor cortices emerges as a promising and quantifiable candidate marker of trait somatic anxiety.

## INTRODUCTION

Anxiety disorders are among the most prevalent psychiatric disorders worldwide [1, 2]. Furthermore, undiagnosed or subclinical anxiety is fairly ubiquitous in the general population and is characterized by impairment in diverse areas of life [3]. The search for the neurological bases of anxiety and anxiety disorders has provided many important insights, yet a comprehensive, translatable understanding has remained elusive. This is reflected in relatively low recovery rate [4] and high prevalence of relapse [5, 6] in anxiety disorders. This warrants further research and, possibly, new approaches.

Research into neurobiology of anxiety primarily focuses on its cognitive aspects. The classical neurocognitive model of anxiety proposes the disruption of the amygdala-prefrontal circuitry in anxiety, which represents deficient recruitment of prefrontal control and amygdala hyper-responsivity to threat [7]. Apart from amygdala-prefrontal circuitry, the cingulo-opercular, fronto-parietal, ventral attention, and default mode networks are among other major networks implicated in the core cognitive mechanisms of anxiety, such as appraising and regulating emotional salience, deranged cognitive control, and so on [8, 9, 10, 11].

However, anxiety is an embodied phenomenon, which is known to cause alterations in several sensorimotor functions including exteroception, interoception and motor functioning. A significant number of individuals with anxiety disorders consistently display symptoms of heightened sensitivity to external stimuli. There appears to be significant epidemiological overlap between anxiety and Sensory Processing Sensitivity (SPS) [12, 13, 14, 15] - a condition characterized by greater sensitivity to subtle stimuli [16, 17]. Several interoceptive symptoms, such as elevated heartbeat perception, palpitation, difficulty breathing, and feeling tense show strong association with anxiety [18, 19, 20, 21, 22, 23]. Elevated muscle tension is among the most consistent physiological findings in chronic anxiety disorders and reduction of muscle tension is a major component of many behavioral therapeutic approaches to alleviate anxiety [24]. Motor restless and hyperactivity are other prominent motor features of anxiety.

Although there are a few neuroimaging studies that examined (or reported) sensorimotor changes in anxiety, our understanding of sensory and motor functions of brain is undergoing a paradigm shift. Motivated by the growing appreciation that top-down/backward information flow in brain (i.e., from functionally higher to lower areas) plays crucial roles in perception (and action) [25, 26, 27], we investigated the effective connectivity among hierarchically organized sensorimotor regions and its association with (trait) anxiety. Here, effective connectivity refers to the influence that one neural system exerts over another, either at a synaptic or population level [28]. In contrast to data-driven approaches (e.g., whole-brain functional connectivity analyses), we adopted a model-based approach informed by empirical knowledge about the functional architectures of sensorimotor networks: namely exteroceptive, interoceptive, and motor networks. For exteroception, we selected lateral frontal pole (FP1)-the terminal relay station for exteroceptive sensory information [29]-and primary visual (V1), auditory (A1), and somatosensory (SSC) cortices. For interoception, we chose anterior and posterior insula (AI and PI) based on known role of the insula in interoception and a posterior to anterior hierarchical organization in insula [8, 30]. Finally for motor regions, we chose supplementary motor area (SMA) and primary motor cortex (MC). SMA is responsible for planning complex movements of the contralateral extremities and is postulated to occupy a higher level of hierarchy in the motor cortex.

Anxiety is known to increase stimulus-driven attention [31], is associated with hypervigilance [32], and psychomotor agitation – states where top-down cortical information flow is supposed to be enhanced. Moreover, as discussed above, anxiety is known to enhance sensorimotor processing in all three sub-domains: exteroception, interoception, and proprioception. Thus, we hypothesized that—with increasing anxiety scores—top-down effective connectivity in three networks will be enhanced. We analyse changes in connectivity associated with both cognitive and somatic aspects of anxiety. Here, somatic represents body (Greek sōmatikos = of the body), as distinct from the mind. The body communicates with the internal and external milieu through sensory perception (exteroceptive, interoceptive and proprioceptive) and action (motor and autonomic). The theoretical motivation for the current study borrows from enactive and embodied treatments that can be seen in the light of interoceptive inference [32, 33, 34, 35, 36]. For example, under embodied cognition hypothesis, “ ‘understanding’ is sensory and motor simulation” [37]. The same holds in the context of mental health: somatic therapy “focuses on resolving the symptoms of chronic stress and post-traumatic stress …… by directing the client’s attention to internal sensations, both visceral (interoception) and musculoskeletal (proprioception and kinesthesis), rather than primarily cognitive or emotional experiences” [38]. The bilateral involvement of interoception and exteroception—two major subdomains of sensory perception and proprioception—underwrites our hypothesis that sensory and motor networks are jointly implicated in somatic anxiety. Specifically, we expected to find correlates of somatic anxiety in sensorimotor hierarchies because many theoretical accounts of anxiety and stress refer to changes in the sensitivity of neuronal populations to afferents: for example, in predictive coding accounts failures of sensory attenuation or selective attention are often cast in terms of aberrant precision or synaptic gain control: e.g., Ainley et al. [39], Barrett and Simmons [40], Gu and FitzGerald [41], and Seth [42].

## MATERIALS AND METHODS

Our main objective was to test the hypothesis of anxiety related enhancement of top-down effective connectivity in sensory and motor networks. Resting state fMRI data from a subset of participants in “WU-Minn HCP Data - 1200 Subjects” dataset available at ConnectomeDB platform (https://db.humanconnectome.org/app/template/Login.vm) were used for that purpose. We used Dynamic Causal Modeling to estimate effective connectivity. A schematic illustration of analysis steps are provided in figure 1.

**Figure 1:**
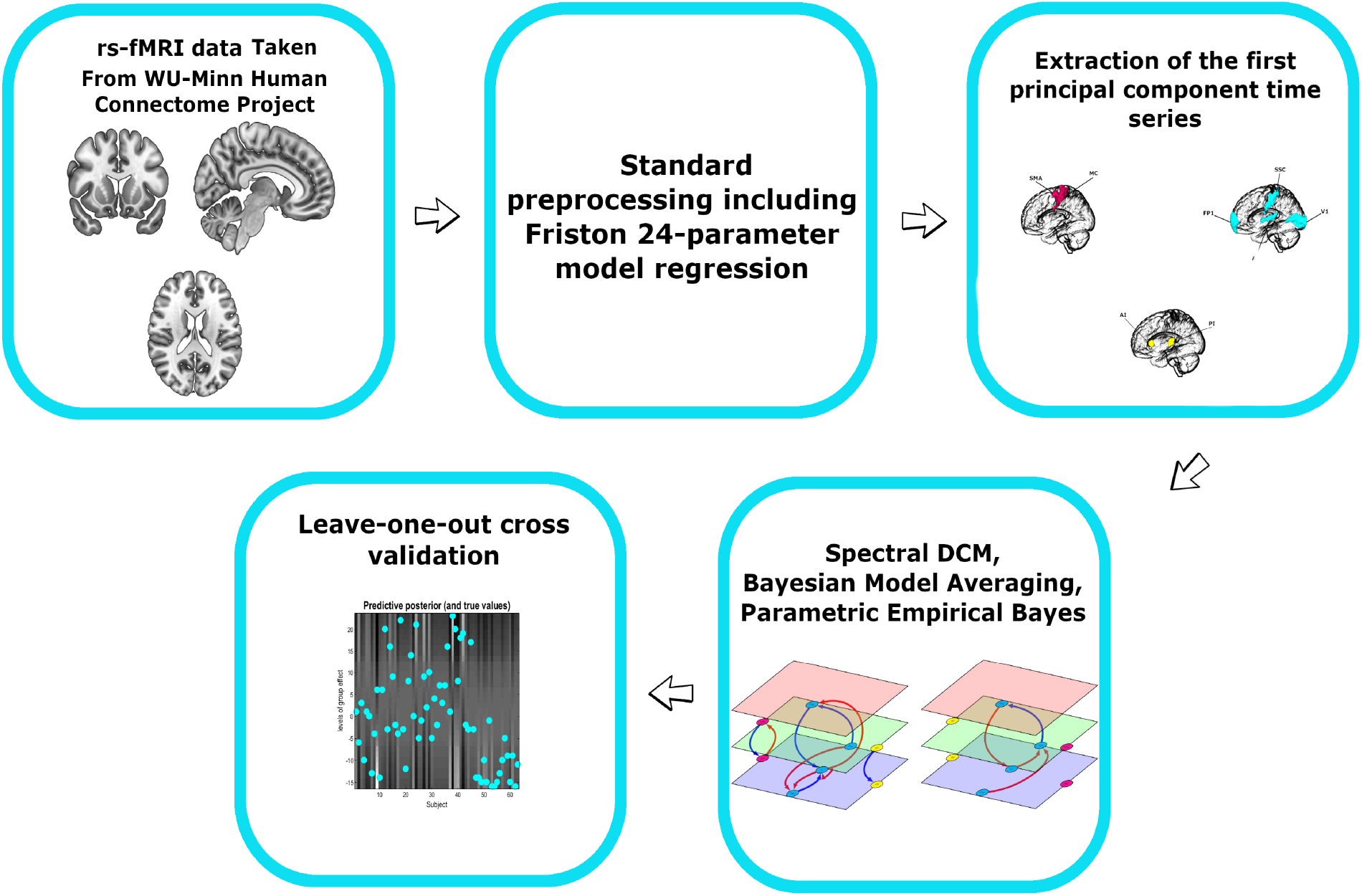
Pipeline of analysis

### Selection of participants and psychometric scores

Among the 1200 participants data available in the dataset, 87 individuals without psychometric scores or MRI scans were eliminated. We also eliminated all the particiapnts with sadness survey scores above 40 which left us with 238 participants. It is well established that depression and anxiety are highly comorbid. Furthermore, a recent study by Ray et al. [43] demonstrated depression associated changes in effective connectivity in sensory and motor cortices. The sadness survey in the NIH toolbox is a self-report measure which assesses sadness using a Computer-Adaptive Test (CAT). According to the NIH toolbox, a sadness score below 40 is considered in the minimal range. By eliminating participants with high sadness score, we controlled for influence of depression on effective connectivity.

Our main psychological measure of interest in this study was the Fear-Somatic Arousal (FSA) survey: unadjusted scale score. The FSA test is a 6-item fixed-length form used in the age group of 18-85 to assess the somatic symptoms of anxiety [44, 45, 46, 47]. The unadjusted scale score has a mean of 50 and standard deviation of 10. We plotted the FSA scores of 238 participants to ascertain the distribution of FSA scores (supplementary figure 6). We found 74 individuals with a score of 40.1. All the scores above 40.1 were distributed over the range of 45-70. A possible reason for this peculiar finding might be some approximation in how the scores are calculated. Since subsequent analysis steps used summary statistics of these scores (e.g., mean, standard deviation), we removed those 74 participants, with the same score, from our analysis. Our final analysis considered 164 participants (81 males and 83 females, mean age 28,7 ± 3.48 years, range 22-35 years).

In addition to FSA, we also included Fear-Affect (FA) Survey: Unadjusted Scale Score in our analysis. This self-report measure assesses fear and anxious misery for ages 18-85, using a CAT format. Overall, FA reflects the cognitive aspects of anxiety.

### Brain image acquisition

All participants were scanned using a Siemens 3T scanner housed at Washington University in St Louis. Scans were taken using a standard 32-channel Siemens receive head coil and a “body” transmission coil. The rsfMRI data were of approximately 15 minutes duration with eyes open and relaxed fixation on a projected bright crosshair on a dark background (in a dark room).

The structural and functional MRI images were acquired using the following parameters [48]:

Structural MRI: T1w 3D magnetization-prepared rapid acquisition with gradient echo (MPRAGE), Field of View: 224 × 224 mm, TR= 2400 ms, TE= 2.14 ms, Flip Angle= 8°, Voxel size= 0.7 mm isotropic. T2W 3D T2-SPACE, Field of View: 224 × 224 mm, TR=3200 ms, TE=565 ms, Flip Angle= variable, Voxel size= 0.7 mm isotropic.

Functional MRI: Gradient echo Echo Planar Imaging (EPI) sequence, Field of View: 208 × 180 mm (*RO × PE*), TR= 720 ms, TE= 33.1 ms, Flip Angle= 52°, Voxel size: 2.0 mm isotropic, 72 slices.

### Pre-processing

We used the minimally pre-processed structural and functional MRI data provided by the Human Connectome Project (WU-Minn HCP 1200 Subjects Data Release) and obtained through the database at http://db.humanconnectome.org. In brief, structural data underwent gradient nonlinearity and b0 distortion correction, co-registration between the T1-weighted and T2-weighted images, bias field correction by capitalizing on the inverse intensities from the T1- and T2-weighting and, finally, registration of the subject’s native structural volume space to MNI space. We used the functional data processed by the fMRIVolume functional pipeline. FMRIVolume removes spatial distortions, realigns volumes to compensate for subject motion, registers the fMRI data to the structural, reduces the bias field, normalizes the 4D image to a global mean, and masks the data with the final brain mask. Details about the preprocessing pipeline could be found in [48]. After downloading these minimally preprocessed data, the first five functional images were removed to allow the magnetization field to stabilize to a steady state and the remaining images were denoised by regressing out several nuisance signals such as head motion (using the Friston-24 head motion parameters) and cerebrospinal fluid and white matter signals using the SPM12 V771 toolbox (Statistical Parametric Mapping, http://www.fil.ion.ucl.ac.uk/spm).

### Selection of ROIs and extraction of time series

The Regions of interest selected for our analysis were the lateral frontal pole (FP1), primary auditory cortex (A1), primary visual cortex (V1) and somatosensory cortex (SSC) for exteroception network, anterior and posterior insulae (AI & PI) for interoceptive network and primary motor cortex (MC) and supplementary motor area (SMA) for motor network. The regions of interest for each network are depicted in figure 2. The ROIs were defined using masks taken from the SPM Anatomy toolbox [49]. To extract time series for subsequent DCM analysis, we took the first principal components of the time series from all voxels included in the masks and adjusted the data for “effects of interest” (i.e., mean-correcting the time series).

**Figure 2:**
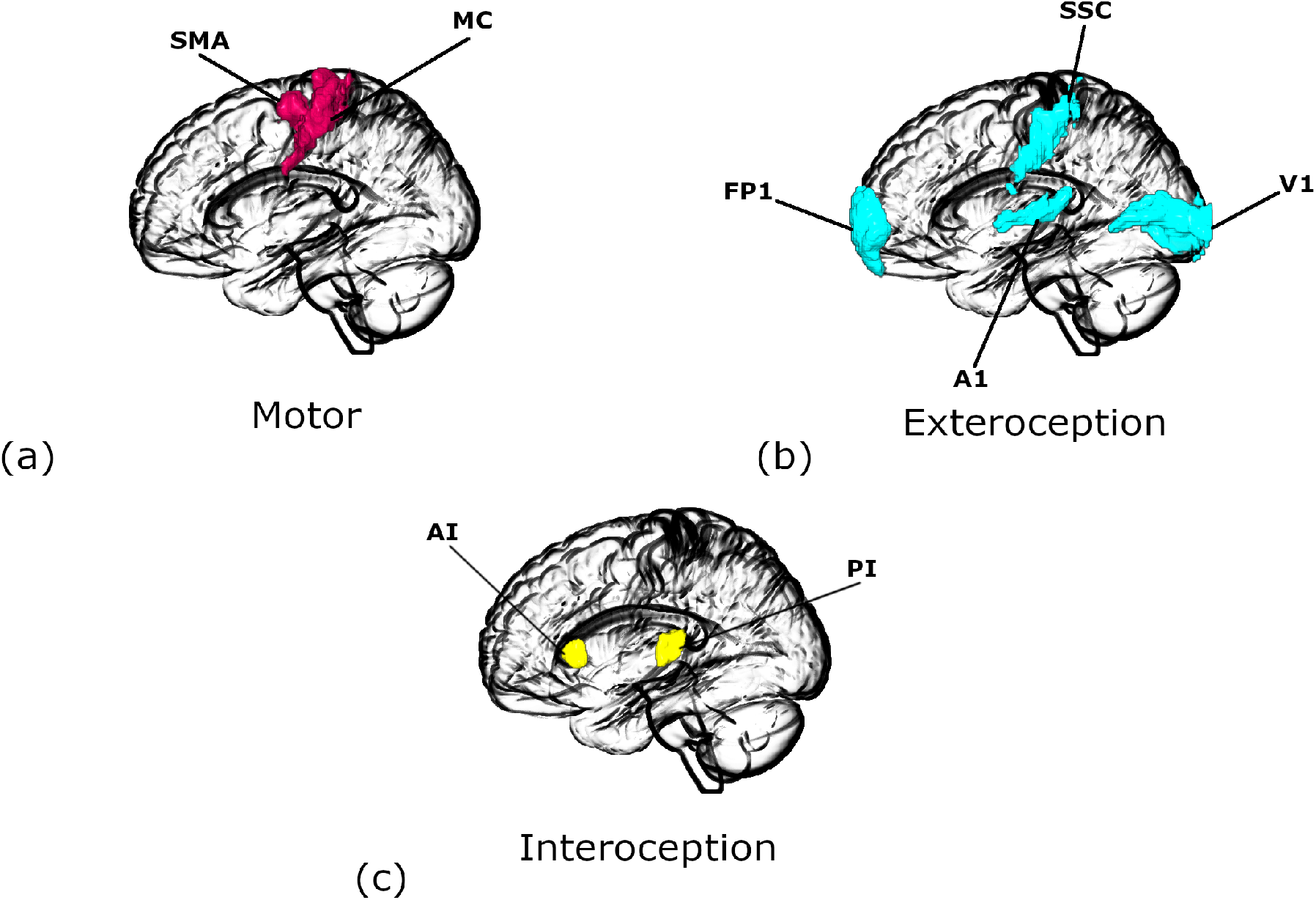
Regions of interest for different networks: (a) Motor. (b) Exteroceptive, and (c) Interoceptive networks. FP1: lateral frontal pole, A1: primary auditory cortex, V1: primary visual cortex, SSC: primary somatosensory cortex, AI: anterior insula, PI: posterior insula, MC: primary motor cortex. SMA: supplementary motor area. The images were created using MRIcroGL (https://www.nitrc.org/projects/mricrogl/).

### Dynamic Causal Modelling and Parametric Empirical Bayes

We used the Spectral DCM approach using DCM12.5 as implemented in SPM12 v7771 (http://www.fil.ion.ucl.ac.uk/spm) to estimate the effective connectivity within each network. Spectral DCM is a computationally efficient form of Dynamic Causal Modeling for estimating effective connectivity from resting state timeseries (as summarized in terms of their cross spectral density).

Dynamic Causal Modeling is a well-established approach to estimate the causal architecture (effective connectivity) of distributed neuronal responses from observed BOLD (Blood-Oxygen-Level-Dependent) signals recorded from fMRI. It is primarily based on two equations. First, the neuronal state equation models the change of a neuronal state-vector in time, depending on directed connectivity within a distributed set of regions that, in the context of DCM for cross spectral density, are subject to endogenous fluctuations, whose spectrum is estimated. Second, an empirically validated hemodynamic model that describes the transformation of neuronal state into a BOLD response. The underlying neural state equation is written as follows:

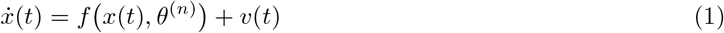

The function *f* is the neural model (i.e., a description of neuronal dynamics), 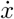 is the rate of change of the neural states *x, θ^n^* is the unknown connectivity parameters (i.e., intrinsic effective connectivity) and v(t) represents a stochastic process that models the endogenous neural fluctuations, which drive the resting state.

The hemodynamic model equation is used to translate the transformation of neuronal state into a BOLD response:

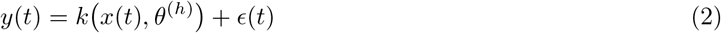

Here, the function *k* specifies the biophysical processes that transform neural activity *x* into the BOLD response with parameters *θ*^(*h*)^ plus the observation noise *ϵ*.

Spectral DCM offers a computationally efficient inversion of the pursuing model of resting state fMRI. Spectral DCM simplifies the estimation of generative model by fitting data features into the frequency domain (i.e., using Fourier transforms) instead of the original BOLD time series as employed in the DCM of evoked induced responses. By switching to second order statistics (i.e., complex cross-spectra), spectral DCM circumvents the problem of estimating time varying fluctuations in neuronal states and estimates their spectra, which is time invariant. In other words, the problem of estimating hidden neuronal states disappears and is replaced by the problem of estimating their correlation functions of time or spectral densities over frequencies (and observation noise) that are much easier to parameterize and estimate. For this purpose, a scale free (power law) form for the endogenous and error fluctuations is used [50] and is written as follows:

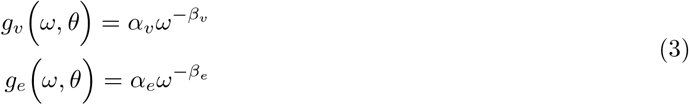

Here, {*α, β*} ⊂ *θ* are the parameters controlling the amplitudes and exponents of the spectral density of these random effects. A standard Bayesian model inversion (i.e., Variational Laplace) is used to infer the parameters of the models from the observed signal i.e., the parameters of the fluctuations and the effective connectivity. A detailed mathematical description of spectral DCM can be found in [27] and [51].

First level (within-subject) analyses involved estimating fully connected models (i.e., between all nodes plus self-connections) for each subject for each kind of network (motor, exteroception, and interoception) in both hemispheres (right and left). In other words, we specified a subgraph or DCM for each modality, with a well-defined lower (i.e., sensory) and higher (i.e., secondary or association) cortical node and estimated the between node (extrinsic) and within-node (intrinsic) effective connectivity for each subgraph. This was repeated for both hemispheres, giving six DCMs for each subject. Our particular interest was in the backward extrinsic connectivity—and whether there were any systematic effects of anxiety on these directed connections, either within or between modalities over subjects. We performed a diagnostic test for each DCM to assess the convergence of model inversion. The accuracy of model inversion was determined by looking at the average percentage variance-explained by DCM model estimation when fitted to the observed (cross spectral) data.

Second level (between-subject) analysis used Parametric Empirical Bayes (PEB): a hierarchical Bayesian model that uses a general linear model (GLM) of subject-specific parameters. The role of PEB is to model differences in individual (within-subject) connections, in relation to subject specific reports of anxiety [52, 53]. One advantage of the PEB framework over the classical tests (e.g., t test) is that it uses the full posterior density over the parameters from each subject’s DCM. That is, both the expected values and the associated uncertainty (i.e., posterior covariance) of the parameters for each connection are taken to the between-subject or group level [54]. The group mean, by default, is the first regressor or covariate in the GLM. FSA scores, FA scores, age, and sex were the further four regressors in this study. FSA and FA scores were mean-centered (across all subjects) to enable the first regressor to be model the group mean of each parameter. Moreover, as FSA and FA scores showed collinearity, we orthogonalized the FA scores with respect to FSA scores using Gram-Schmidt orthogonalization. We plot the FSA, FA, and orthogonalized FA scores in figure 3. We used Bayesian model reduction (BMR) to explore the space of possible models that could potentially explain the resting state data in all subjects. BMR considers candidate models by removing one or more connections from a full or parent model [53]. BMR prunes connection parameters from the full model by scoring each reduced model based on its log model-evidence. The pruning continues until there is no further improvement in model-evidence. The final step accommodated uncertainty over the remaining models using Bayesian Model Averaging (BMA) [55]. BMA takes the parameters of the selected models and averages them in proportion to their model evidence.

**Figure 3:**
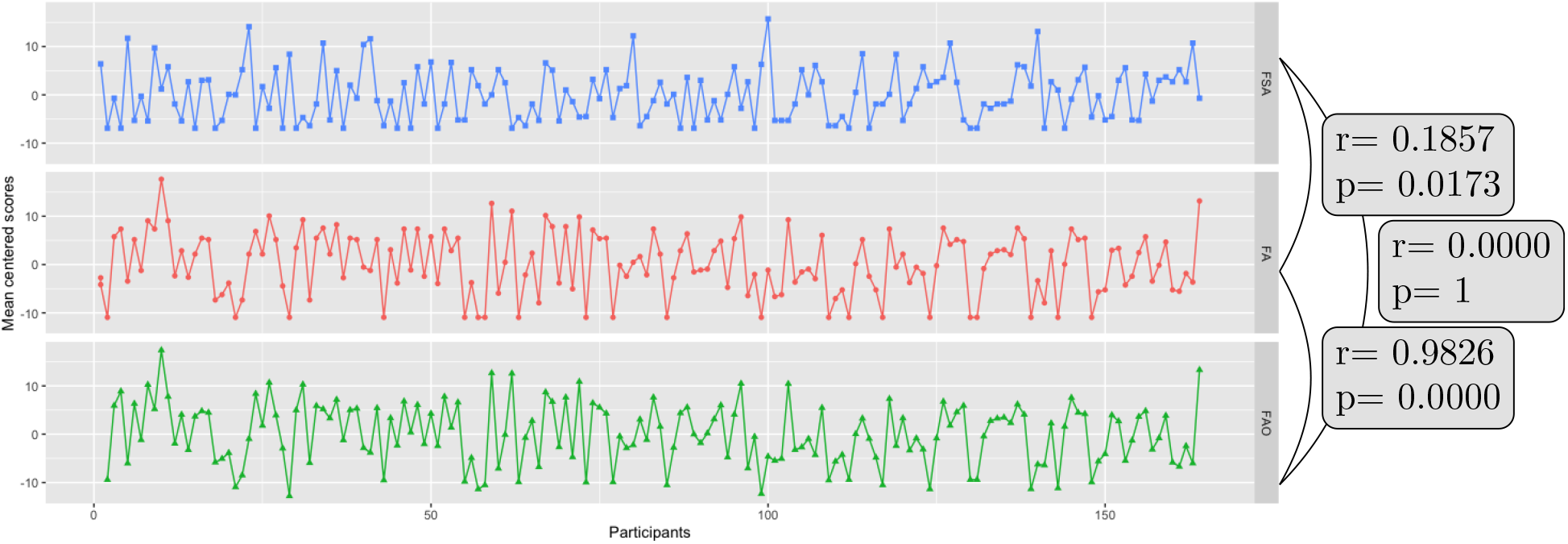
Psychometric scores across participants before and after othogonalization. FSA: fear somatic arousal scores, FA: fear affect scores before orthogonalization, FAO: fear affect scores after orthogonalization. The correlation values between difference scores are also shown.

### Leave-one-out validation analysis

Finally, we tested whether the FSA scores could be predicted based on subject-specific estimates of effective connectivity using a leave-one-out cross validation analysis [56]. To include just those parameters (i.e., connection strengths) that had the greatest evidence of being non-zero, we thresholded the BMA to focus on parameters that had a 95% posterior probability of being present. A participant was left out and a parametric empirical Bayesian model was inverted to predict the score of the left-out subject, based on the specific connections chosen. The Pearson’s correlation between the predicted score and the measured was used to summarize the out-of-sample effect size; namely, the degree to which effective connectivity could predict – or could be predicted by – anxiety.

## RESULTS

### Accuracy of DCM model estimation

The inversion of DCM models for individual participants produced excellent results in terms of accuracy (see figure 4). Across participants, the minimum percentage variance-explained by DCM -when fitted to observed (cross spectra) data - were 79.82%, 57.69% and 53.26% for right motor, exteroception and interoception networks respectively. As for the left hemisphere, these values were 77.64%, 44.31% and 48.89% respectively.

**Figure 4:**
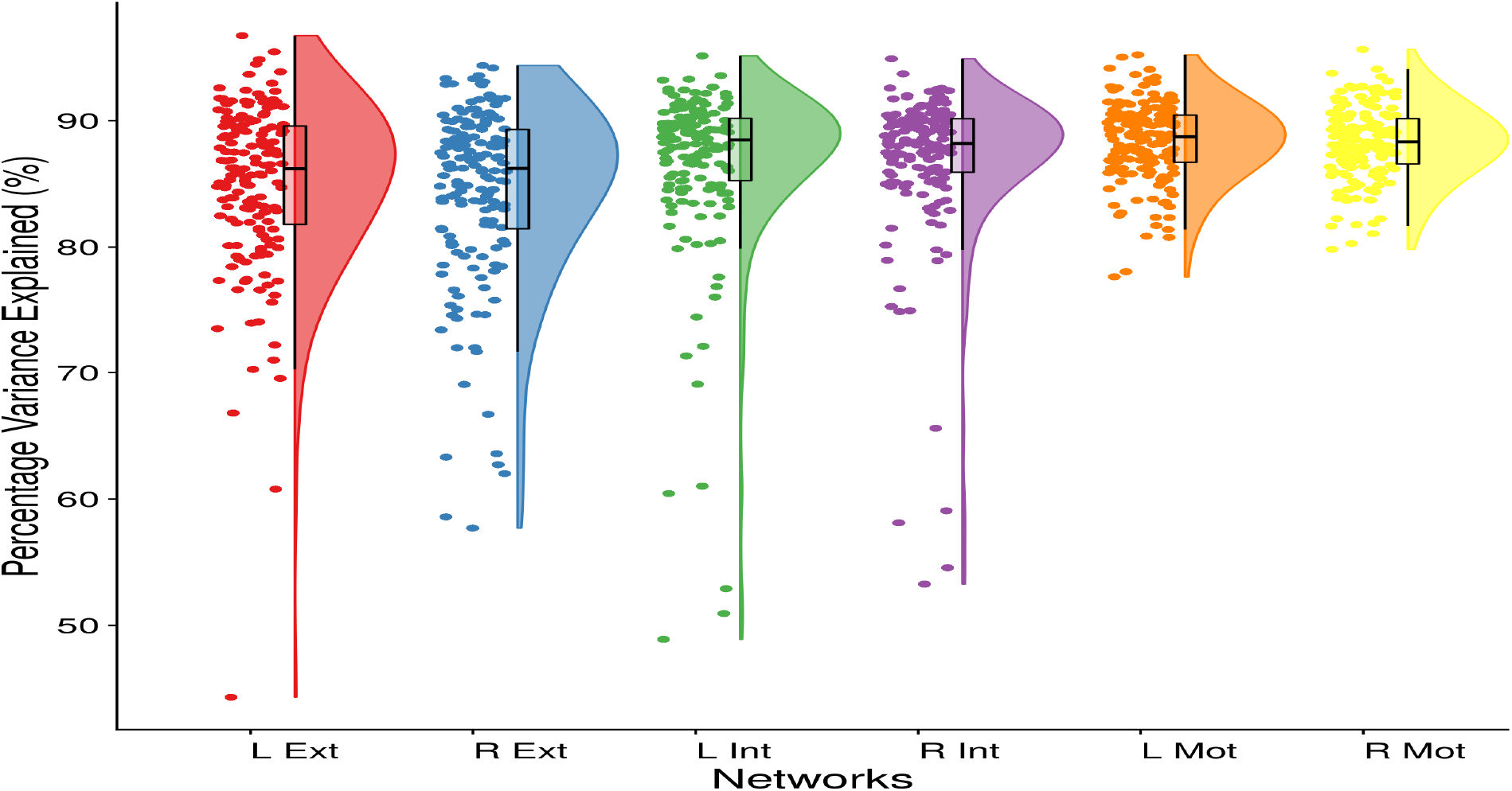
Accuracy of DCM model estimation: Average-percentage explained by our DCM models for the target networks in both hemispheres.

### Effective connectivity

#### Group mean effective connectivity

The results for group mean effective connectivity are shown in figure 5 (a) and (b) and detailed in supplementary figure 1. Among extensive networks of connections in both hemispheres, one noteworthy pattern emerged in the backward effective connectivity. In exteroceptive sensory networks across hemispheres, backward connections were inhibitory. In interoception and motor networks, they were excitatory. Most of the forward connections, in all three networks, were inhibitory in nature.

**Figure 5:**
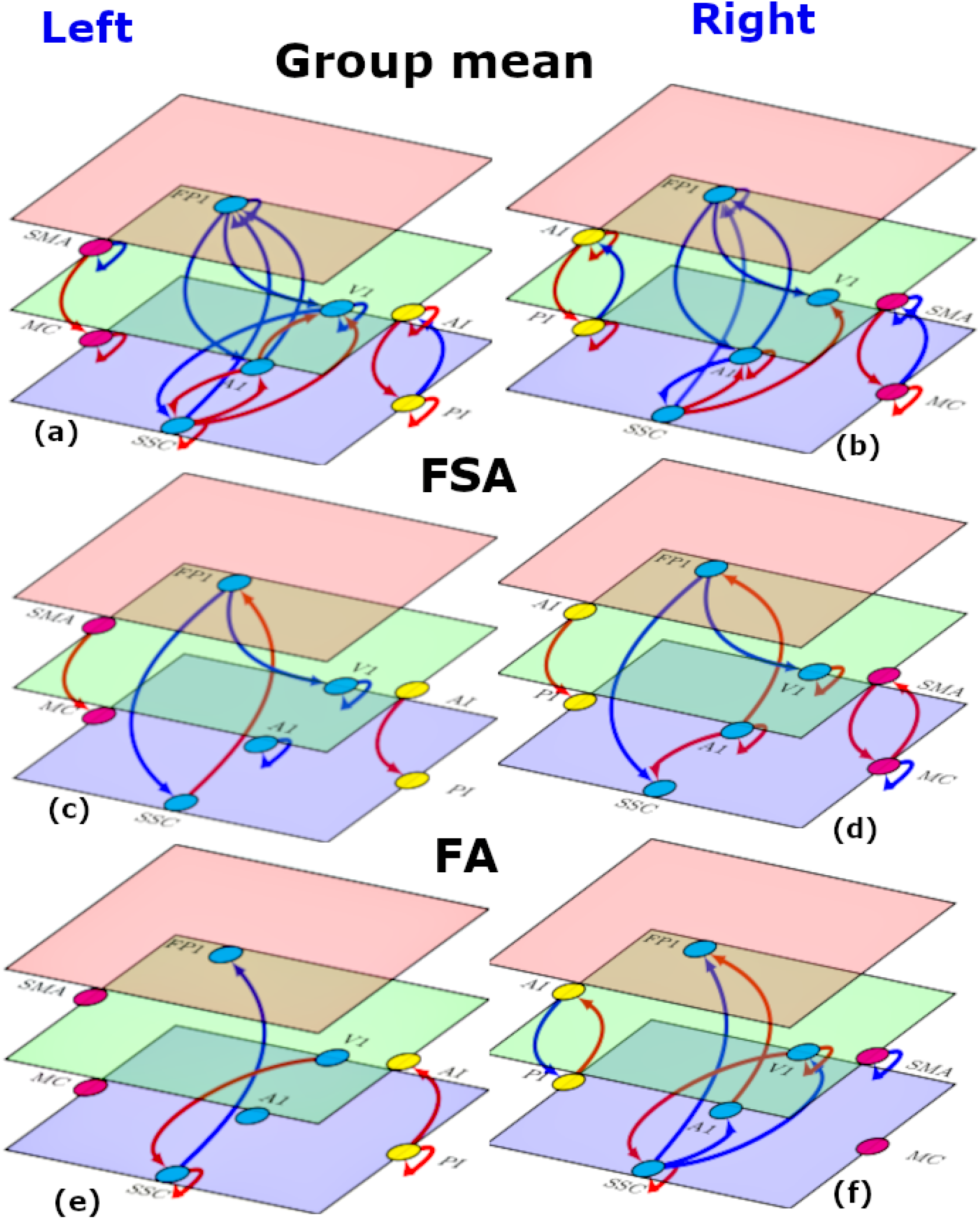
Effective connectivity (left and right hemispheres). (a),(b): Group mean effective connectivity in sensory and motor networks. Arrow colours code nature of connections red, excitatory; blue, inhibitory. (c),(d),(e),(f): Connections showing significant association with FSA (c and d) and FA (e and f) scores in sensory and motor networks. Arrow colours code direction of connectivity changes relative to the group mean: red, increased; blue, decreased. For all subfigures line thickness is kept constant and does not code for the effect size. For the exact values of the estimated connectivity parameters see Appendix A. Nodes are placed in different planes to denote relative position of different nodes in cortical hierarchy. SMA: supplementary motor area, MC: primary motor cortex, FP1: lateral frontal pole, V1: primary visual cortex, A1: primary auditory cortex, SSC: primary somatosensory cortex, AI: anterior insula, PI: posterior insula.

#### Changes in effective connectivity with FSA scores

FSA scores reflect the somatic aspect of trait anxiety. The overall pattern of relative changes in effective connectivity with increasing FSA scores is depicted in figure 5 (c) and (d). The actual values can be found in supplementary figure 2. As with mean connectivity, the FSA associated changes were most consistent in (extrinsic or between region) backward connections across both hemispheres. In general, all three kinds of network showed enhanced top-down (backward) influences. For example, in exteroceptive cortices, with increasing FSA scores top-down connections became more inhibitory. Conversely, top-down connections in interoceptive and motor networks became more excitatory.

#### Changes in effective connectivity with FA scores

FA scores primarily represent cognitive aspects of anxiety. The connections showing significant association with FA scores are depicted in figure 5 (c) and (d). Supplementary figure 3 contains estimated effective connectivity values. Unlike FSA scores, no enhancement of top-down connectivity was observed in association with FA scores. Rather, right AI to PI backward connectivity was weakened i.e., became inhibitory (it was excitatory at the group mean level). In general, changes with FA scores remained confined to bottom-up and lateral connections.

#### Cross validation

In a leave-one-out cross-validation among all connections showing significant association with FSA scores, five connections were found to predict FSA scores at a significant level of *α* = 0.05 (see Table 1 and figure 6). These connections were: left FP1 to left SSC (corr=0.22, p-value=0.00272),, right FP1 to right V1 (corr=0.17, p-value=0.01682), right A1 to right V1 (corr=0.14, p-value=0.03359), right MC to right MC (self-loop) (corr=0.15, p-value=0.02801), left AI to left PI (corr=0.15, p-value=0.02792). Note that these correlation coefficients are not corrected for multiple comparisons because they are not testing hypotheses – they are summarizing out-of-sample effect sizes, under the most likely model of anxiety-related changes in effective connectivity among subjects.

**Figure 6:**
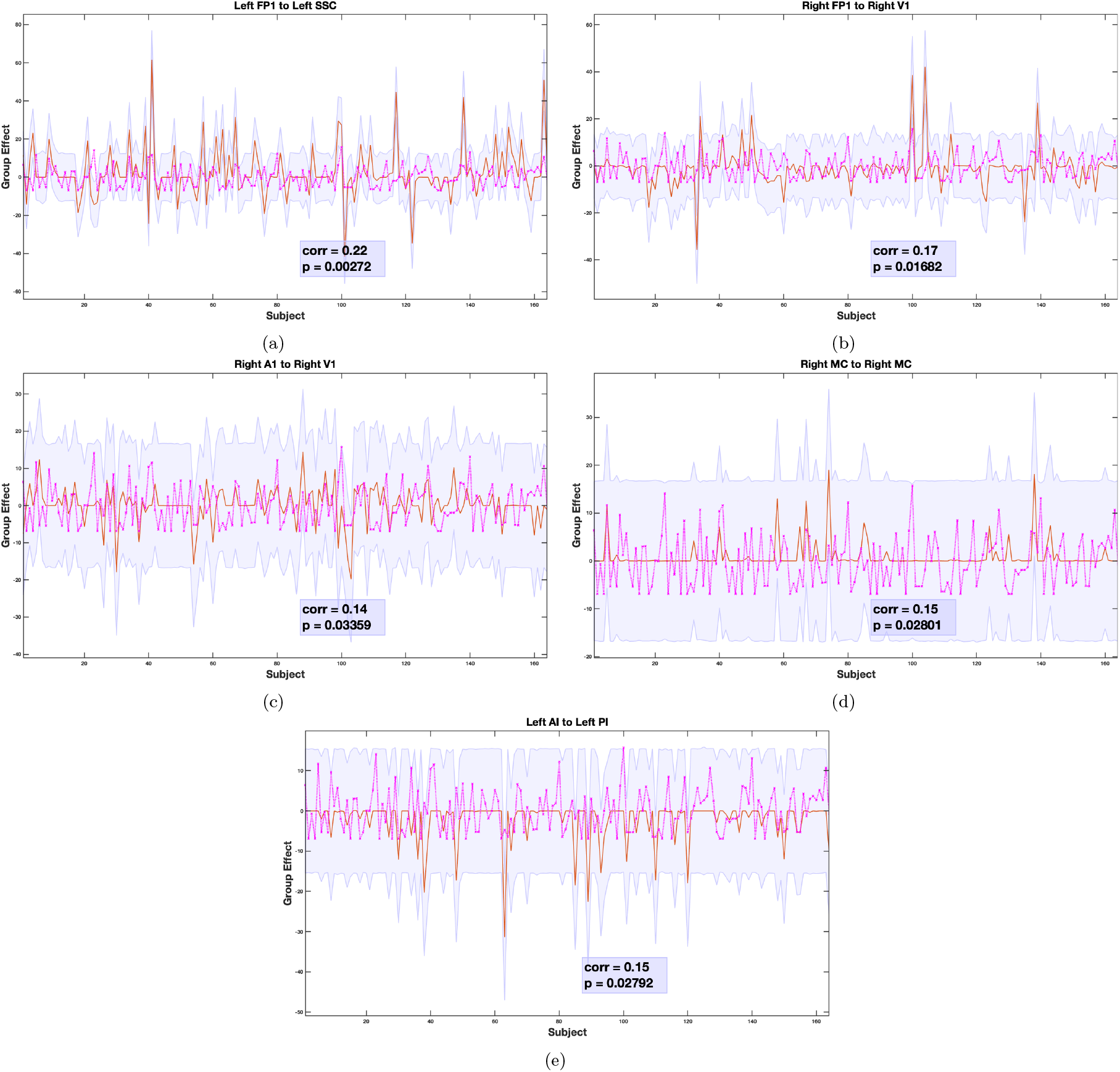
Leave one out cross validation analysis: the out-of-samples estimate of FSA score for each subject (orange line) with 90% confidence interval (shaded area). The purple line represents the actual FSA score. Correlation values between actual and estimated FSA scores are shown in the legend.

**Table 1:** Leave-one-out cross validation.

| Connections | Correlation | p value |
| --- | --- | --- |
| **lFP1 → lSSC** | **0.22** | **0.00272\*** |
| lFP1 → lV1 | 0.08 | 0.14339 |
| lSSC → lFP1 | 0.04 | 0.28516 |
| lV1 → lV1 | 0.06 | 0.23341 |
| rFP1 → rSSC | -0.02 | 0.61198 |
| **rFP1 → rV1** | **0.17** | **0.01682\*** |
| rA1 → rFP1 | 0.12 | 0.06161 |
| rA1 → rSSC | 0.09 | 0.12070 |
| **rA1 → rV1** | **0.14** | **0.03359\*** |
| rA1 → rA1 | 0.13 | 0.05109 |
| lSMA → lMC | -0.06 | 0.78262 |
| rSMA → rMC | 0.05 | 0.24576 |
| rMC → rSMA | 0.12 | 0.5771 |
| **rMC → rMC** | **0.15** | **0.02801\*** |
| **lAI → lPI** | **0.15** | **0.02792\*** |

## DISCUSSION

The most striking result of our study was found in the group-averaged backward (top-down) effective connectivity in sensory and motor cortices that showed consistent patterns across hemispheres and consistent changes with FSA scores. The backward effective connections in exteroceptive sensory networks were inhibitory by nature whereas they were excitatory in bilateral interoceptive and motor networks. With increased FSA scores (but not with increased FA scores), this pattern was accentuated in all three networks. In other words, top-down inhibition in exteroceptive network and top-down excitation in interoceptive and motor networks were associated with increased report of somatic anxiety. In leave-one-out cross validation analyses, five connections were found to have a sufficiently large effect size to predict whether somebody has a high or a low FSA score.

There are a few neuroimaging studies that have examined or reported functional/effective connectivity changes in sensory and motor cortices as biomarkers for anxiety [57, 58, 59, 60, 61, 62]. In contrast to these extant studies, our work is motivated by a novel understanding of neural mechanisms underlying sensory perception. There is a growing consensus that perception is not a passive ‘bottom-up’ mechanism of progressive abstraction from sensory input. Rather, recurrent information flow between hierarchically organized brain regions plays a crucial role in perception. This underwrites most current theorizing about functional brain architecture. The most prominent is the theory of predictive coding [25, 26, 27] that has also been extended to motor domain [63, 64]. The implicit neuronal message passing within cortical hierarchy motivated us to investigate effective connectivity among hierarchically organized sensorimotor regions and its association with (trait) anxiety.

At the group level, the most consistent finding in the current study concerns the top-down average effective connectivity in different sensory and motor networks. Across participants and hemispheres, top-down connections are inhibitory in exteroceptive networks and excitatory in interoceptive and motor networks. The pattern in exteroceptive network is consistent with the role of top-down predictions explaining away prediction errors at lower levels, as proposed by the predictive coding framework [27, 65]. Although long-range connections in the brain are excitatory (i.e., glutamatergic), predictive coding proposes that backward connections may preferentially target inhibitory interneurons in superficial and deep layers to evince an overall decrease in neuronal message passing [27, 65, 66]. In predictive coding, this is often read as ‘explaining away’ prediction errors at lower levels in sensory cortical hierarchies so that only those incoming stimuli that deviate from prediction (i.e., prediction errors) ascend the hierarchy to revise presentations at higher levels [65, 66, 67]. However, the completely opposite pattern was observed in interoceptive and motor networks. Descending excitatory connections in interoceptive and motor systems may reflect the increased precision that is thought to mediate attention and sensory attenuation [68, 69, 70]. In this instance, there can be an explaining away of certain prediction errors, while their precision may be increased, resulting in an overall excitatory drive. In terms of synaptic physiology, this can be read as an increase in postsynaptic gain or cortical excitability.

Moreover, a complementary explanation exists for top-down excitatory connectivity in motor networks. It could be a reflection of the fact that ascending prediction errors in the executive motor system may play a small role – because these prediction errors are thought to be resolved through cortical spinal reflexes, i.e., through action [64]. Put simply, in sensory hierarchies exteroceptive prediction errors are caused by bottom-up sensory input, which are resolved by (inhibitory) top-down predictions. Conversely, in motor hierarchies, prediction errors are generated by (excitatory) top-down proprioceptive predictions, which are resolved by motor reflexes at the level of the spinal-cord.

In line with the marked consistency of the patterns of top-down effective connectivity, the changes in backward effective connectivity—with the extent of somatic arousal—also showed a consistent pattern and fit comfortably with phenomenology of anxiety syndromes. With increasing somatic anxiety scores, the patterns found in top-down connections at the group level are accentuated in all three networks. In other words, connections from lateral frontal pole to primary exteroceptive cortices become more inhibitory, whereas connections from supplementary motor area to primary motor cortex—and from anterior to posterior insula become more excitatory with increasing somatic anxiety. Enhancement of top-down processing in exteroceptive networks is in line with the hypervigilance and heightened sensitivity to external stimuli in anxiety and increased Sensory Processing Sensitivity (SPS) [16, 17] in several anxiety disorders. The enhancement in the interoceptive network is consistent with elevated heartbeat perception, difficulty breathing, feeling tense [18, 19, 20, 21, 22, 23] and other interoceptive symptoms in anxiety. Elevated muscle tension, motor restless and hyperactivity are some of the prominent features of anxiety [24] that might well be reflected in the increased top-down motor network effective connectivity.

In this context it is noteworthy that descending predictions of precision play an important role in active inference accounts of several psychiatric conditions – in which synaptic pathophysiology and psychopathology can be accounted for by a failure of sensory attenuation; namely, the attenuation or suspension of the precision of sensory prediction errors. This failure of attention and attenuation has been used to explain several conditions, including autism, schizophrenia, Parkinson’s disease and depression [71, 72, 73, 74, 75]. The present work suggests that the disinhibition—which underlies a failure of sensory attenuation—may also characterize trait somatic anxiety.

In sharp contrast to FSA scores, FA scores did not show association with increased descending effective connectivity among regions in exteroceptive, interoceptive and motor networks. This finding is in line with-and furnishes the neurobiological basis for-distinction between cognitive and somatic aspects of anxiety [76, 77, 78, 79, 80, 81, 82, 83, 84, 85, 86]. In this work, we effectively define somatic anxiety in relation to cognitive anxiety. Cognitive anxiety has been broadly described as the “negative expectations, worries, and concerns about oneself, the situation at hand, and potential consequences”. In contradistinction, somatic anxiety has been associated with “the perception of one’s physiological arousal” [87]. Cognitive anxiety has been frequently associated with expressions like “fear of the worst happening”, “terrified”, and “fear of losing control” whereas somatic anxiety emphasises the interoceptive and autonomic sequalae of embodied states; including shortness of breath, pounding heart, dizziness, sweating, numbness, unsteadiness, feeling hot, and a feeling of choking [88, 89, 90, 91, 92, 93]. While cognitive and somatic symptoms interact with one another [94], they have been postulated to be distinctive aspects of the anxiety process, reflected in differences in antecedence, phenomenology, behavioral relevance, and therapeutic response [76, 77]. For example, threat of electric shock has been shown to have its primary influence on somatic anxiety, whereas social or performance evaluation tends to have a stronger eliciting effect on cognitive anxiety [78, 79]. Similarly, cognitive anxiety symptoms have been shown to respond effectively to cognitively-oriented approaches such as cognitive restructuring or processing. On the other hand, somatic symptoms have been shown to respond to physiologically-based approaches, including biofeedback and relaxation[80, 81]. Finally, cognitive and somatic anxiety have differential influences on performance. In several studies in professional athletes, cognitive anxiety is unrelated to motor performance [82] or has a negative relationship [83], whereas somatic anxiety is accompanied by improved perceptuo-motor performance [84] or demonstrates an inverted-U relationship with motor performance: such that performance initially improved and then deteriorated with increasing somatic anxiety [83]. Note that the finding from the present study does not suggest that individuals with heightened cognitive anxiety cannot show increased backward effective connectivity as cognitive and somatic anxiety are often coexistent. The results, however, do suggest that the enhanced top-down cortical processing in sensorimotor networks reflects somatic, rather than cognitive, aspect of anxiety.

The model comparison discussed above furnishes clear evidence for changes in a number of connections that underwrite anxiety, as scored with the FSA scores. One might now ask whether these changes can be used to predict somatic arousal in individuals. In other words, are the underlying effect sizes sufficiently large to anticipate whether somebody has a high or a low FSA score? This question goes beyond whether there is evidence for an association and addresses the utility of connectivity phenotyping for precision medicine. One can estimate out of sample effect sizes using cross validation under a parametric empirical Bayesian scheme [53]. In this analysis, we withhold a particular participant and ask whether one could have predicted the FSA score given the effective connectivity estimates from that subject. In the current analysis, five individual connections from three networks showed a significant out-of-sample correlation with FSA score. This suggests that a nontrivial amount of variance in the FSA score could be explained by effective connectivity. It is also important to note that three out of five connections are backward connections, further highlighting the importance of top-down processing in somatic anxiety.

A note on our choice of network nodes. As we were primarily interested in quantifying top-down or descending connectivity in the cortical hierarchy, we selected primary sensory/motor cortices and an accompanying “higher” region for each network. Thus bilateral primary motor, visual, auditory, somatosensory cortices and posterior insula were selected as lower nodes. It should be pointed out here that posterior insula is widely considered as the primary interoceptive cortex [95, 40, 96]. For motor networks, SMA was chosen as the higher node. SMA is responsible for planning (rather than directly executing) complex movements of the contralateral extremities [97, 98] and is thus posited to occupy a higher level in the functional hierarchy than the primary motor cortex. For interoception, we chose the anterior insula based on a posterior to anterior hierarchical organization in the insula [99, 100]. For exteroceptive networks, the higher region ought to be situated at a higher level in the functional organization of the cortex compared to three lower regions: the primary visual, auditory, and somatosensory cortices. Several tracing, lesion, and physiological studies suggest that visual, auditory, and somatosensory processing pathways converge at different regions of VLPFC [101] and DLPFC [102]. We therefore chose the lateral frontal pole as representative of a higher node. Empirical studies [103, 104] support a posterior to anterior sensory representational hierarchy in the prefrontal cortex and place the lateral frontal pole one level higher than both DL and VL PFC in the cortical hierarchy.

Findings from the current study should be considered within the context of certain limitations. Although our study sample was large for neuroimaging measures-and we undertook steps like cross-validation to ensure the generalizability of our findings-replication in a different sample would be an important next step. Secondly, in the present work, we investigated the association of effective connectivity with subthreshold anxiety in a large-scale dataset. None of our participants reported a diagnosis of anxiety disorder. We will consider testing for the association of pathological anxiety with effective connectivity, in sensory and motor networks, in future work.

The insights from the present work have significant translatable potential. Further research should be conducted to investigate the effectiveness of therapeutic interventions like neurofeedback and relaxation techniques in reverting the top-down effective connectivity and physical symptoms of somatic anxiety. Another interesting approach is emerging noninvasive brain stimulation techniques like Transcranial Magnetic Stimulation (TMS). Recent research has established that TMS can modulate cortico-cortical connectivity in specific neural circuits [105, 106, 107]. Specific brain regions in the sensory or motor networks could be stimulated and their effect on the somatic anxiety could be studied using state-of-the-art techniques that are currently available. This may have clinical implications for pathological or trait anxiety in the general population and in specific contexts like sports performance.

In summary, our results advance our mechanistic understanding of the pathophysiology that underwrites somatic anxiety. Traditional neuroimaging accounts of anxiety have centered around the cognitive aspects of anxiety at the expense of bodily symptoms. The present work establishes anxiety as an embodied phenomenon by demonstrating that enhanced top-down effective connectivity in sensory and motor cortices affords a promising neural signature of trait anxiety. It also establishes the generalizability and predictive validity of this novel marker– and may portend a new avenue of research into the neural underpinnings and therapeutic treatments of anxiety.

## DATA AND CODE AVAILABILITY

Our analysis code is available on GitHub (https://github.com/dipanjan-neuroscience/anxiety2021).

Imaging data are available on connectomeDB platform of Human Connectome Project (https://db.humanconnectome.org/app/template/Login.vm).

## ACKNOWLEDGEMENTS

This research was supported by the Basque Government through the BERC 2018-2021 program, by the Spanish Ministry of Science, Innovation, and Universities (BCBL Severo Ochoa excellence accreditationSEV-2015-0490 and BCAM Severo Ochoa accreditation SEV-2017-0718), the Spanish Ministry of Economy and Competitiveness (Ramon y Cajal Fellowship, RYC-2017-21845) and the project MTM2017-82379-R(AEI/FEDER,UE) (principal investigator: Dr. Maria Xose Rodriguez, BCAM). K.J.F. was funded by a Wellcome Trust Principal Research Fellowship (Reference 088 130/Z/09/Z). We are thankful to the Neuroscience Gateway (NSG) (83) for the computational resources provided. For the purpose of Open Access, the authors have applied a CC BY public copyright licence to this manuscript.

## AUTHOR CONTRIBUTIONS

D.R., and M.D. conceived the present project. I.B., D.R., and M.D. performed data analysis. C.C.G, K.J.F, and D.R supervised the project. I.B., D.R., and M.D. wrote the manuscript. C.C.G and K.J.F edited the manuscript.

## Supplementary figures

**Supplementary Figure 1:**
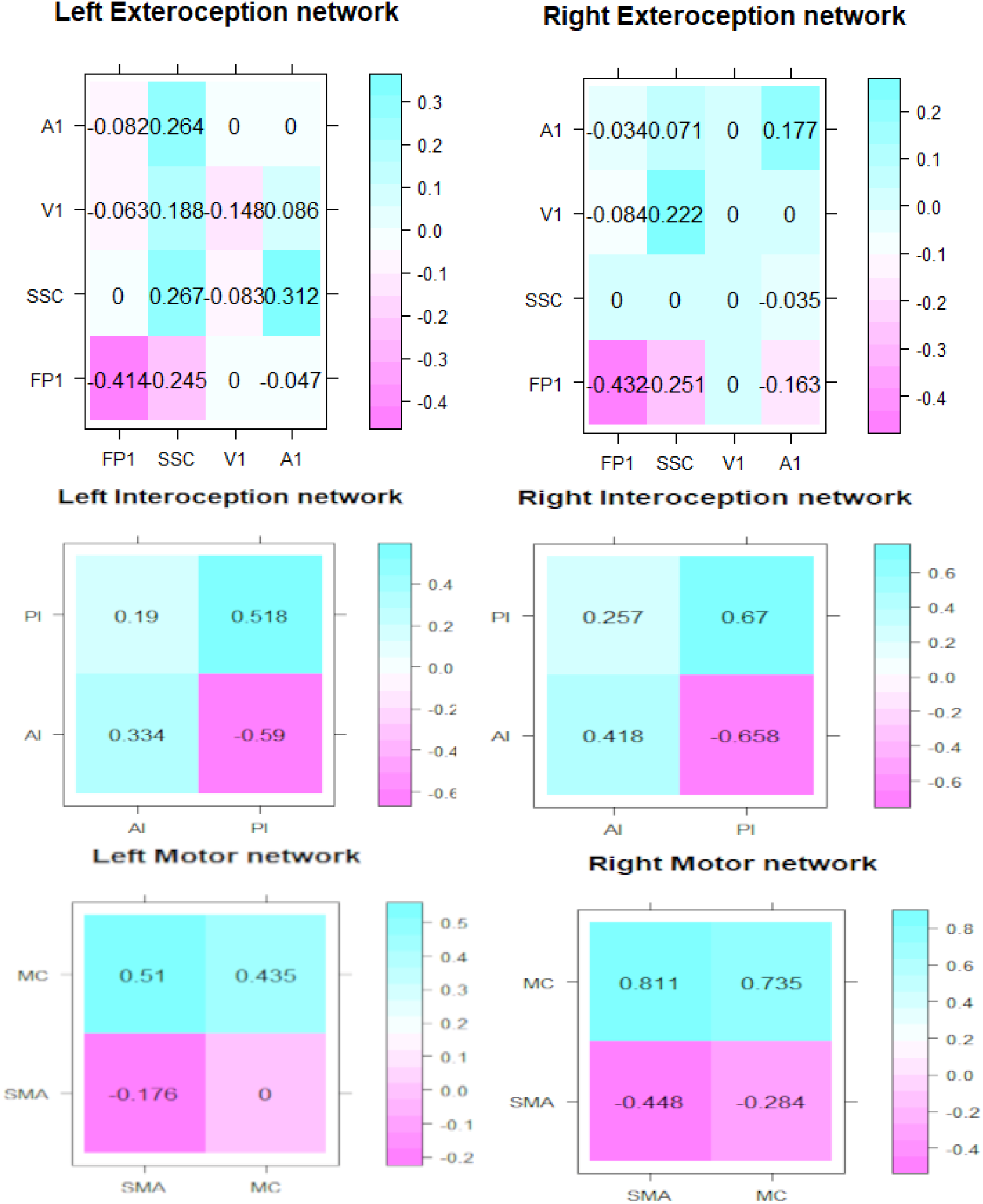
Mean connectivity values (columns: from, rows:to)

**Supplementary Figure 2:**
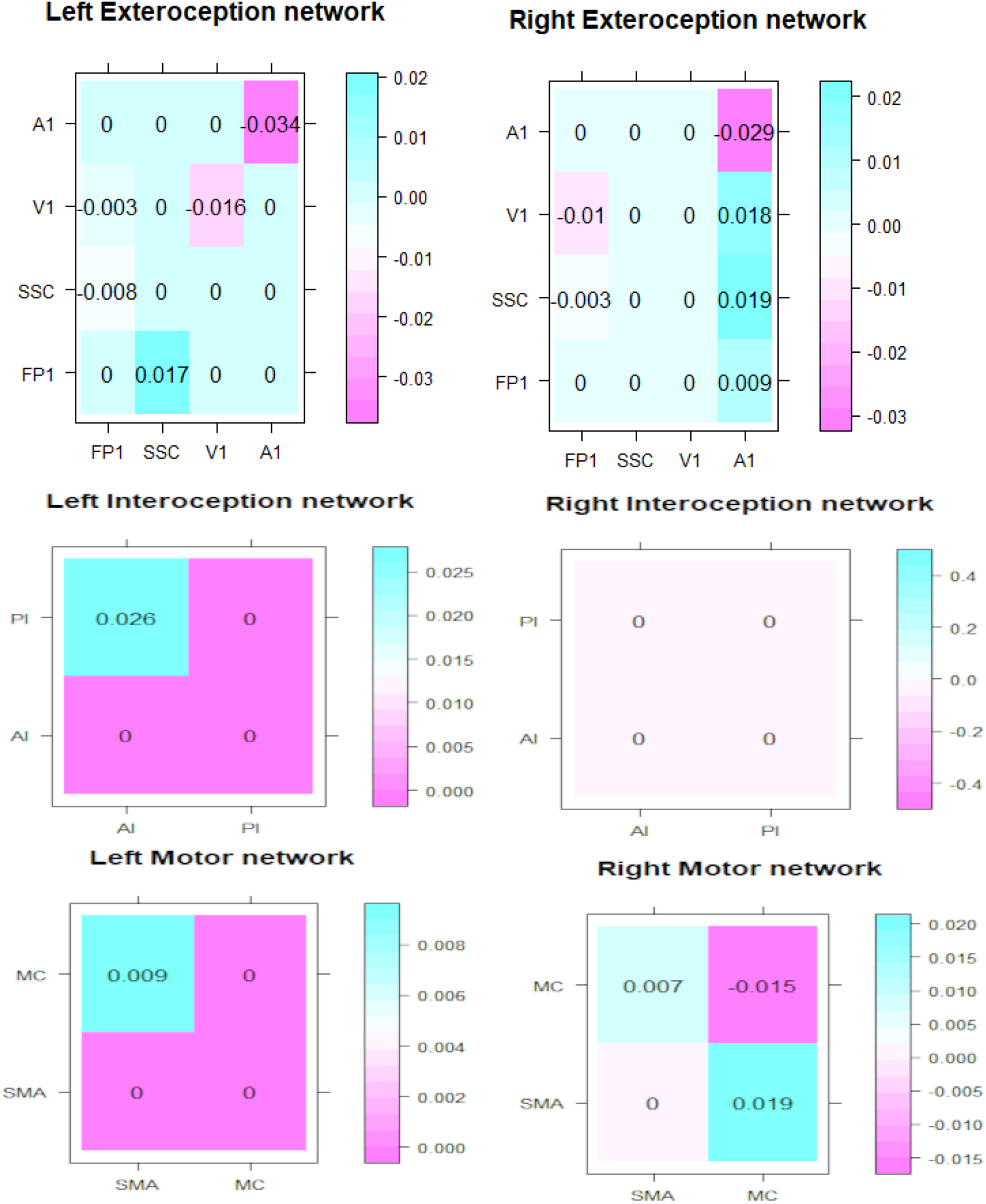
FSA connectivity values (columns: from, rows:to)

**Supplementary Figure 3:**
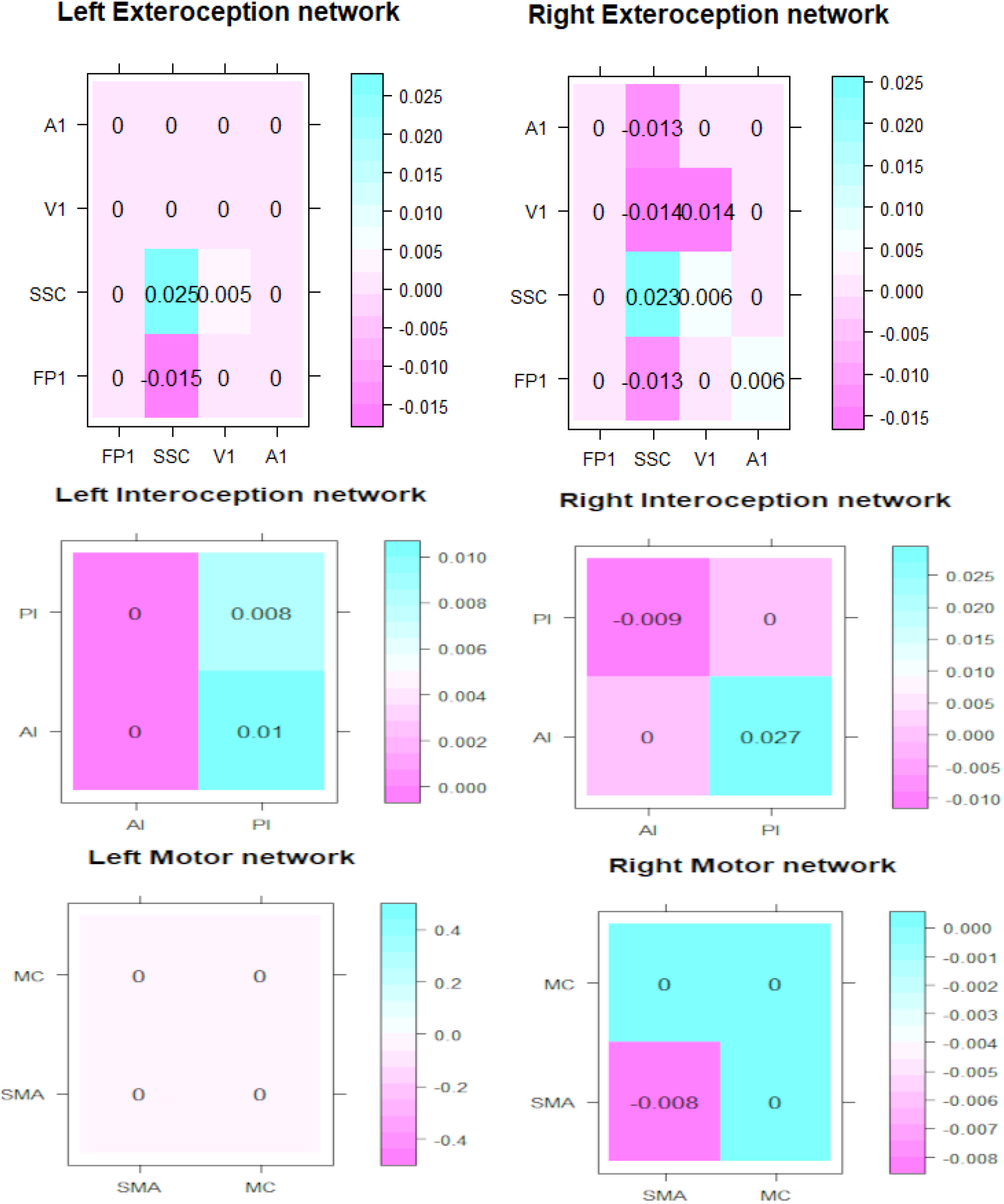
FA connectivity values (columns: from, rows:to)

**Supplementary Figure 4:**
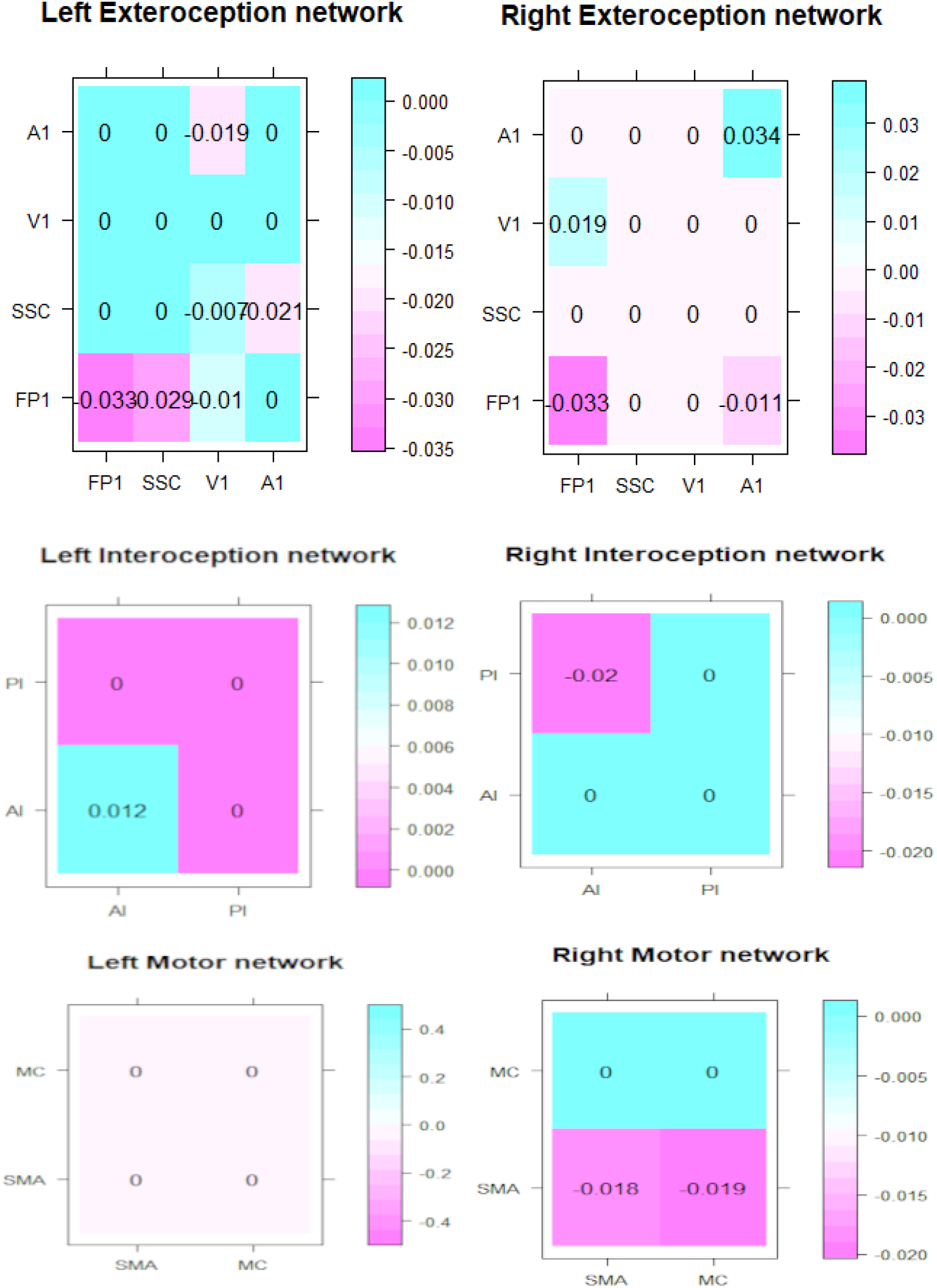
Age connectivity values (columns: from, rows:to)

**Supplementary Figure 5:**
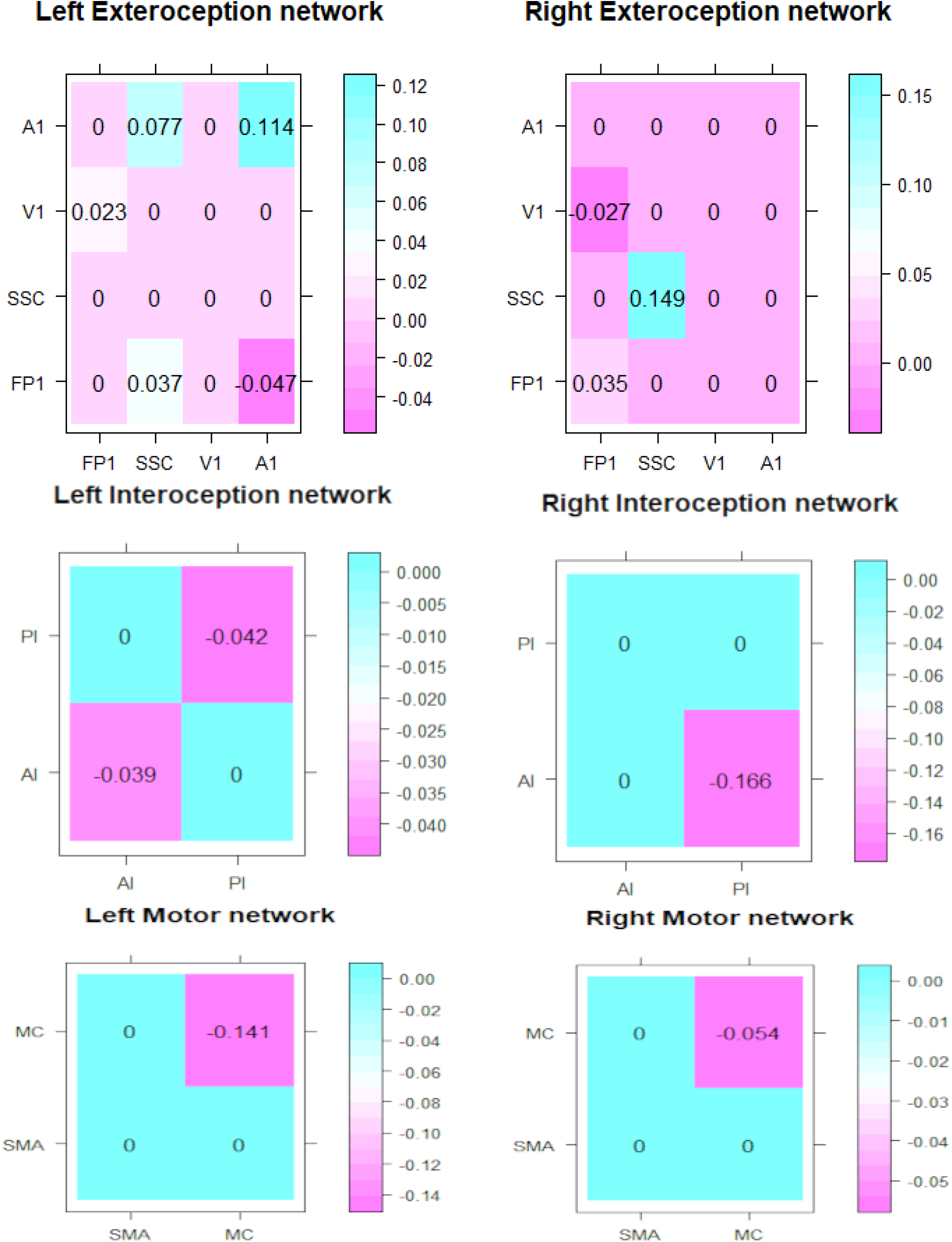
Sex connectivity values (columns: from, rows:to)

**Supplementary Figure 6:**
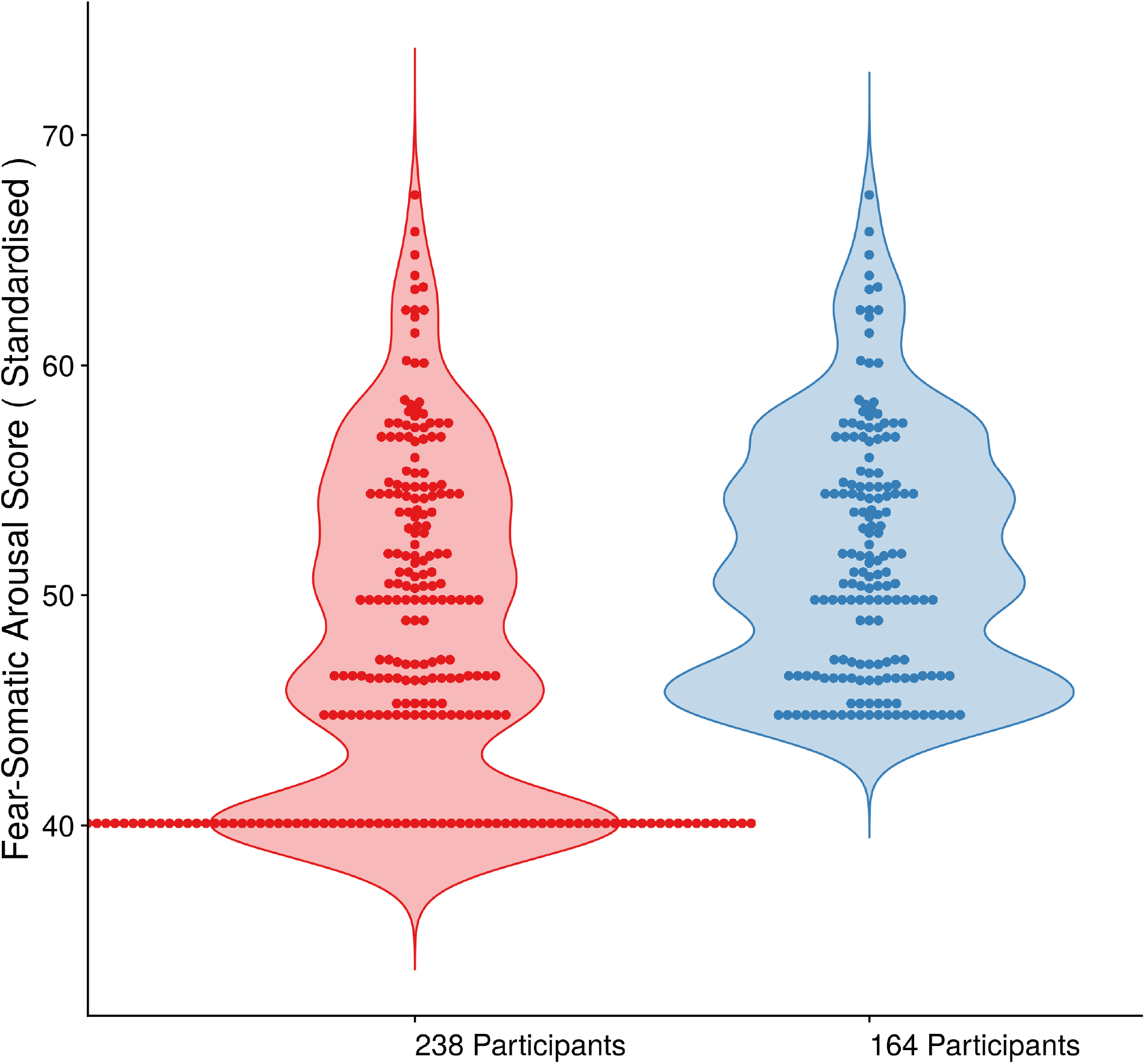
Frequency distribution of Fear-Somatic Arousal Score before and after elimination of 74 participants

